# GRASP: Gene-relation adaptive soft prompt for scalable and generalizable gene network inference with large language models

**DOI:** 10.1101/2025.10.20.683485

**Authors:** Yuqiang Feng, Kaiwen Deng, Yuanfang Guan

## Abstract

Gene networks (GNs) encode diverse molecular relationships and are central to interpreting cellular function and disease. The heterogeneity of interaction types has led to computational methods specialized for particular network contexts. Large language models (LLMs) offer a unified, language-based formulation of GN inference by leveraging biological knowledge from large-scale text corpora, yet their effectiveness remains sensitive to prompt design. Here, we introduce Gene-Relation Adaptive Soft Prompt (GRASP), a parameter-efficient and trainable framework that conditions inference on each gene pair through only three virtual tokens. Using factorized gene-specific and relation-aware components, GRASP learns to map each pair’s biological context into compact soft prompts that combine pair-specific signals with shared interaction patterns. Across diverse GN inference tasks, GRASP consistently outperforms alternative prompting strategies. It also shows a stronger ability to recover unannotated interactions from synthetic negative sets, suggesting its capacity to identify biologically meaningful relationships beyond existing databases. Together, these results establish GRASP as a scalable and generalizable prompting framework for LLM-based GN inference.

## Introduction

Cells are governed by complex molecular interactions that collectively shape cellular states. These interactions can be abstracted as gene networks (GNs), graph-based representations of genes that coordinate cellular processes^1^. Understanding how genes interact within such networks is fundamental to deciphering biological processes and disease mechanisms^2–5^. GNs encompass diverse forms of biological relationships, including protein-protein interaction (PPI) networks that capture physical contacts among proteins, gene regulatory networks (GRNs) that describe directed transcriptional control, and phosphorylation networks that represent kinase-substrate modification events^6–8^.

The diversity of GNs has motivated computational approaches based on different data modalities. Sequence-based models learn interaction-relevant features from protein amino acid sequences to predict PPIs^9–11^. Gene expression-based methods infer regulatory relationships from co-expression patterns and perturbation responses^12-13^. Graph-based approaches leverage network topology and graph learning techniques to capture higher-order relational structure^14-15^.

Recent advances in large language models (LLMs) offer a complementary paradigm by capitalizing on biological knowledge internalized during pretraining on scientific text. In this setting, GN inference can be formulated as a language-based prediction problem:given a pair of genes, an LLM can draw on its biological knowledge to assess whether a relationship is likely to exist. This formulation enables a unified treatment of different GNs without requiring modality-specific feature engineering.

However, LLM-based inference in the biomedical domain is highly sensitive to prompt design^16-17^. This sensitivity poses a practical challenge for GN inference, where a prompting strategy must generalize across highly diverse gene pairs while remaining scalable to millions of candidate interactions. Fixed prompts provide identical context for all pairs, failing to capture the heterogeneity of gene functions and interactions. Naively appending descriptive text to the prompt can be counterproductive, as verbose or irrelevant context may obscure rather than reinforce the relational signal.

These observations highlight a key challenge: how to condition LLM-based GN inference on gene interaction context in a robust and parameter-efficient manner. Soft prompts provide a promising alternative by introducing learnable virtual tokens that guide inference during fine-tuning^18^. Yet standard soft prompts learn a single shared embedding per task, limiting their capacity to capture instance-level biological variability.

To address this limitation, we present Gene-Relation Adaptive Soft Prompt (GRASP), an instance-adaptive framework that synthesizes a small set of virtual tokens conditioned on each queried gene pair. GRASP captures pair-specific signals through gene-dependent coefficients while leveraging shared interaction patterns through a common prototype basis. By generating only three adaptive virtual tokens per pair, GRASP provides a scalable mechanism to improve LLM-based GN inference across different interaction types. Across large-scale PPI inference, single-cell perturbation benchmarks, and phosphorylation network inference, GRASP consistently outperforms both task-specific soft prompts and fixed prompts augmented with biological gene descriptions.

## Results

### Prompting methods for gene network inference

We applied and benchmark the prompting strategies on two LLMs: Gemma-3-4B-IT^19^ and Llama-3.1-8B-Instruct^20^. To enhance the models’ capacity for capturing biologically meaningful gene relationships, we continue-pretrained (CPT) them on domain-specific corpus comprising titles and abstracts from 6.3 million gene-related PubMed articles^21^ (Fig. 1A). For each architecture, we retained both the original and domain-adapted variants, yielding four backbone configurations for subsequent evaluation.

**Figure 1.**
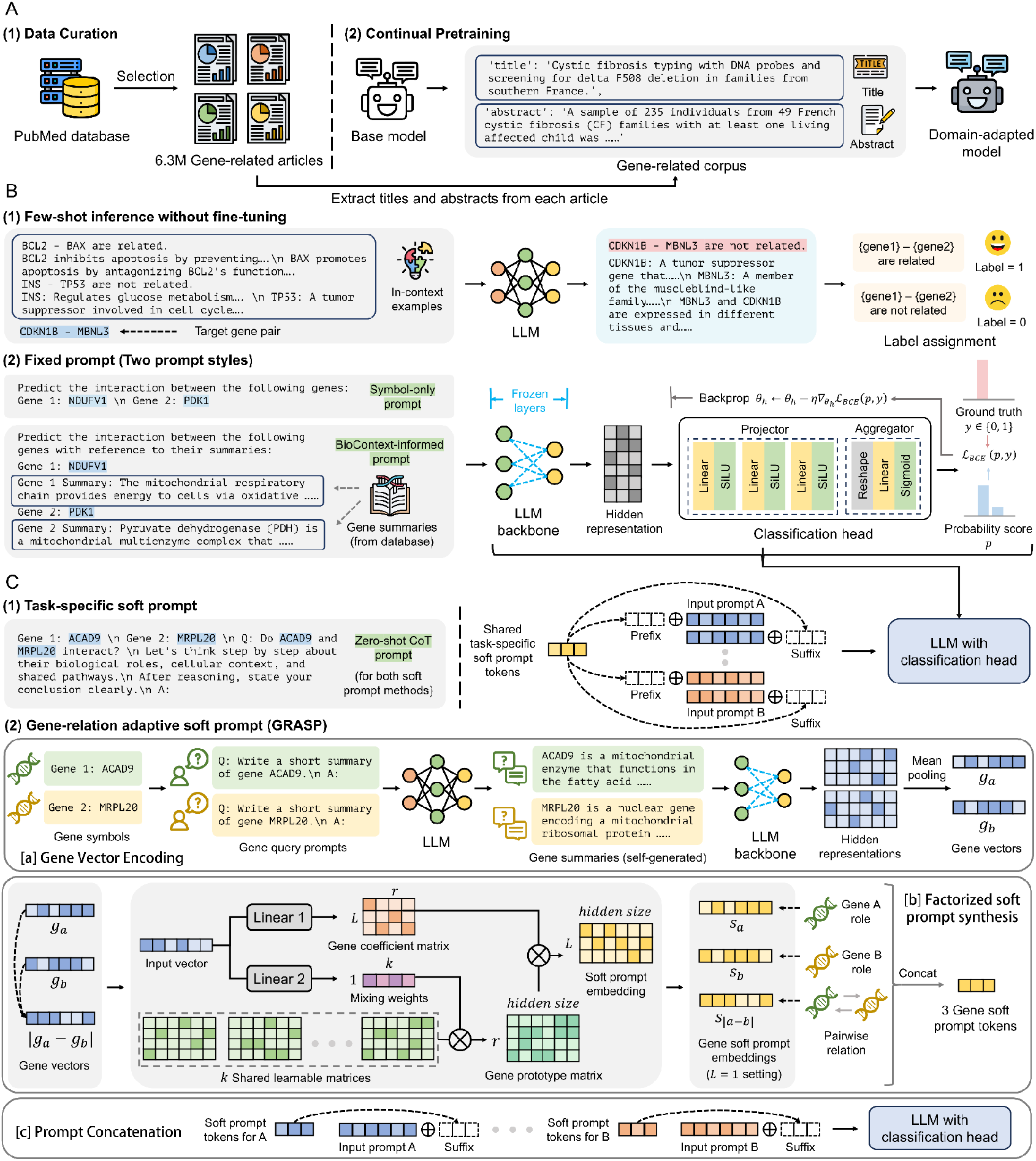
Framework overview. **(A)** Domain-specific continual pretraining. Approximately 6.3 million gene-related articles were curated from PubMed. Titles and abstracts were extracted to form a gene-focused corpus for continual pretraining. **(B)** Baseline methods. Few-shot inference relies on in-context examples. Fixed prompts employ either symbol-only or biocontext-informed templates, followed by a classification head trained on representations from a frozen LLM to predict gene interactions. **(C)** Soft prompt-based methods. Task-specific soft prompts learn a shared set of virtual tokens across gene pairs. GRASP generates instance-adaptive soft prompts via three stages: gene vector encoding, factorized soft prompt synthesis, and prompt concatenation.

Across these configurations, we compared GRASP against several prompting baselines spanning discrete and continuous approaches (Fig. 1B, C). Few-shot inference provides a minimal-effort reference by supplying a small number of labeled gene-pair examples without parameter updates (Fig. 1B(1)). For all other methods, the original language modeling head is replaced with a lightweight classification head, which is fine-tuned while the LLM backbone remains frozen. Under this setting, we evaluated two fixed-prompt designs: a symbol-only template containing only gene symbols, and a biocontext-informed template that additionally incorporates short gene descriptions from curated databases (Fig. 1B(2)). We also implemented a task-specific soft prompt baseline following standard prompt tuning practice^18^, in which a single shared set of learnable virtual tokens is applied uniformly to all gene pairs (Fig. 1C(1)). Both soft prompt methods use the same zero-shot chain-of-thought textual prompt as input context^22^.

Building on the task-specific soft prompt formulation, GRASP extends soft prompting by conditioning the virtual tokens on each input gene pair (Fig. 1C(2)). For each gene, GRASP first generates a concise summary using the backbone LLM and encodes it into a fixed vector via mean pooling of the final-layer hidden states. Given a gene pair, three context vectors are constructed: two for the individual genes and a third encoding their element-wise absolute difference. Each context vector is then mapped to a soft prompt token through a factorized formulation comprising a gene-specific coefficient matrix and a shared prototype matrix—the former captures pair-specific signals via linear projection, while the latter represents common interaction patterns as a weighted combination of learnable bases. This yields exactly three adaptive virtual tokens per gene pair, appended as a suffix to the input prompt.

### GRASP improves large-scale PPI inference with cross-species transferability

We first evaluated GRASP on a large-scale benchmark of human protein-protein interactions (PPIs) comprising 2,127,124 gene pairs, balanced between experimentally validated physical interactions and randomly sampled putative non-interacting pairs. Physical PPIs were obtained from curated resources including BioGRID^23^, MIPS^24^, IntAct^25^, and STRING^26^. We fine-tuned models on this dataset using a standard train-validation-test split: 1,361,359 training pairs (64.0%), 340,340 validation pairs (16.0%), and 425,425 test pairs (20.0%). Each experiment was repeated five times with different random splits, and mean ± SD values were reported. Performance was assessed on 50,000 randomly selected test pairs using precision, recall, accuracy, and ROC-AUC to capture both overall prediction quality and sensitivity-specificity trade-offs.

Across all backbone configurations, GRASP achieved the best precision-recall trade-off, reaching the highest precision at comparable recall levels (Fig. 2A). Few-shot inference yielded the lowest performance across all metrics, while other methods clustered at intermediate levels, clearly separated from GRASP. Numerically, GRASP achieved average relative improvements of 6% in precision and 10% in recall over the baselines (excluding few-shot inference). ROC-AUC analyses corroborated these findings: GRASP achieved 0.923 and 0.931 with Gemma 4B, and 0.925 and 0.937 with Llama 8B before and after domain adaptation, surpassing all other methods (Fig. 2B). Consistent gains in accuracy further support its robustness (Supplementary Fig. 1A).

**Figure 2.**
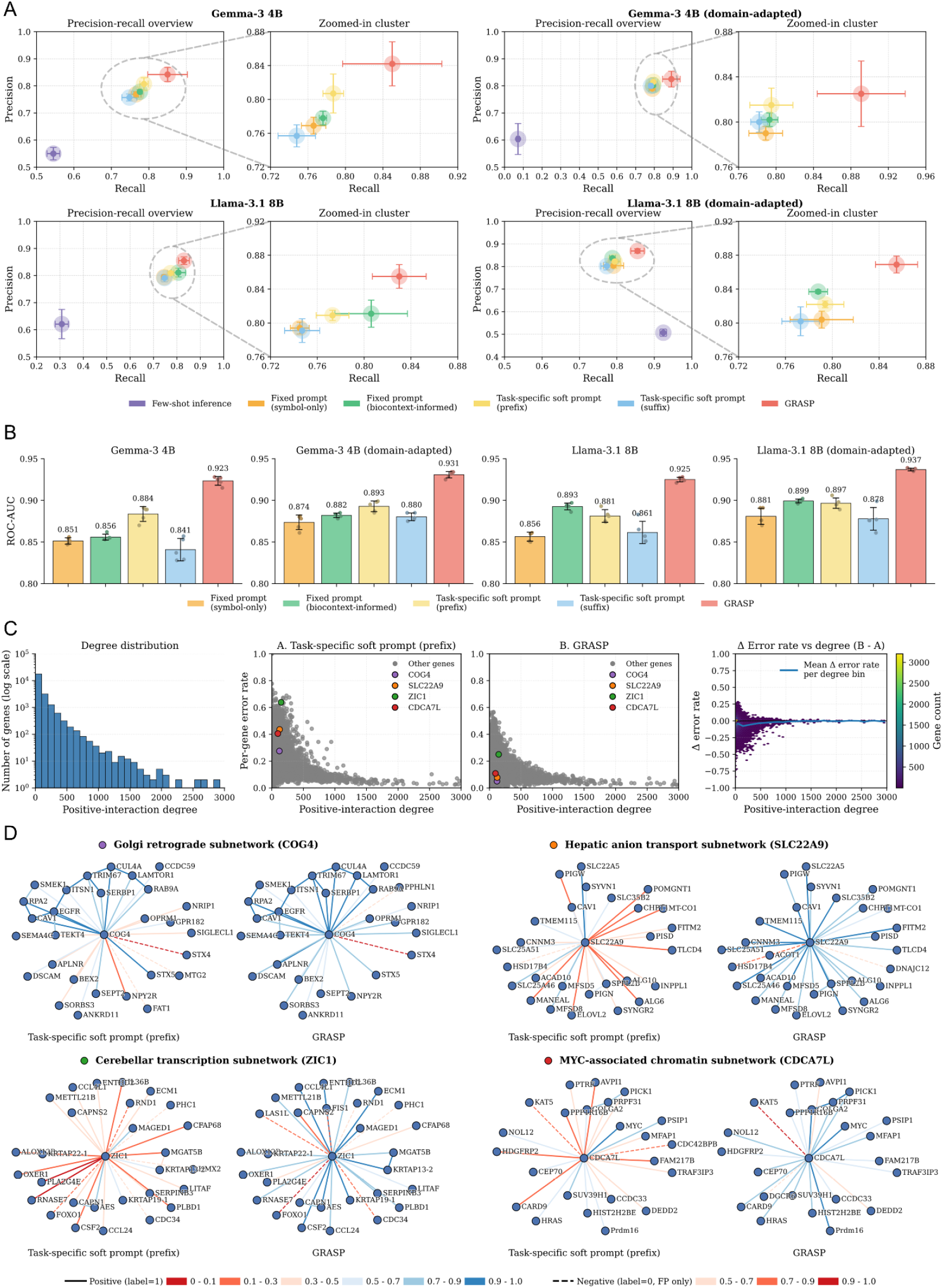
Comparative performance on the human PPI dataset. **(A)** Precision-recall analyses for Gemma-3 4B and Llama-3.1 8B models, with and without domain adaptation. Each panel includes an overview plot (left) and a zoomed-in view of high-performing clusters (right). **(B)** ROC-AUC scores for all prompting methods. **(C)** Per-gene error rate analyses. Left: degree distribution of positive interactions (log scale). Middle: scatter plots of per-gene error rates versus positive-interaction degree. Highlighted genes illustrate differences in error patterns. Right: Δ error rates between task-specific soft prompt (prefix) and GRASP, plotted against positive-interaction degree, with negative values indicating improvements by GRASP. The blue line denotes the mean Δ error rate per bin. **(D)** Subnetwork visualizations of representative genes (COG4, SLC22A9, ZIC1, and CDCA7L). Edge colors indicate prediction outcomes: solid blue, true positives (TPs); solid red, false negatives (FNs); dashed red, false positives (FPs).

To assess whether these improvements were broadly distributed or concentrated among specific gene classes, we examined per-gene error rate distributions along with the gene degrees (number of interactions the gene has involved). GRASP achieved the lowest error rates for high-degree genes and showed even greater relative improvements among moderately connected genes, where other methods exhibited systematic misclassifications (Fig. 2C, Supplementary Fig. 2, 3). Representative subnetwork analyses illustrate this advantage (Fig. 2D). In the COG4 subnetwork, GRASP recovered known retrograde transport partners (NPY2R, BEX2) while reducing false positives; in the ZIC1 subnetwork, it retained transcriptional regulators (RNASE7, OXER1) missed by task-specific soft prompts. These cases demonstrate that GRASP not only improves global metrics but also reconstructs more biologically coherent local network structures.

To further assess cross-species transferability of GRASP, we directly applied models fine-tuned on the human PPI dataset to chicken, cow, and dog PPI datasets derived from the Integrated Interactions Database (IID)^27^, with all pairs overlapping with the human dataset removed. Each dataset was balanced by adding an equal number of randomly sampled non-interacting pairs, resulting in 153,252 pairs for chicken, 165,574 for cow, and 149,960 for dog. Evaluation procedures were identical to those applied in the human PPI setting.

Cross-species transfer proved challenging, with precision-recall performance tightly clustered among methods (Supplementary Fig. 4A). This reduced separation is likely due to the exclusion of gene pairs overlapping with the human training set, leaving predominantly species-specific interactions that are underrepresented in the model’s learned knowledge. GRASP favored higher precision with a modest reduction in recall, yielding comparable F1 scores to baselines. As a threshold-dependent metric, F1 score may not fully capture performance under domain shifts where optimal decision boundaries vary. In contrast, ROC-AUC provided a more robust measure of relative ranking quality, revealing a clearer separation. Under this metric, GRASP achieved the highest AUC in most conditions (Supplementary Fig. 4B). Together, these results indicate that GRASP exhibits stronger cross-species generalization than other prompting methods.

### GRASP achieves strong biological performance on single-cell perturbation benchmarks

To evaluate GRASP in a biologically grounded setting with real perturbation data, we turned to CausalBench^28^, a benchmark built on large-scale Perturb-seq datasets from K562 and RPE1 cell lines. CausalBench supports two complementary evaluation modes: biological evaluation, which quantifies agreement with curated gene-gene interactions, and statistical evaluation, which measures distributional consistency with perturbation-induced expression changes.

We constructed undirected interaction candidates between perturbed and observed genes, and applied models fine-tuned on human PPIs to infer these regulatory relationships in a zero-shot setting. All pairs overlapping with the PPI training data were removed to prevent information leakage. Importantly, LLM-based methods, including GRASP, do not use gene expression profiles but rely solely on text-derived representations. As an expression-based baseline, we included GRNBoost^12^ and evaluated the method under three regimes: purely observational data, 50% interventional data, and 100% interventional data.

On both K562 and RPE1 cell lines, GRASP achieved the highest biological F1 score among all methods (Fig. 3A, 3B, top). Compared with other prompting strategies, GRASP operated at higher precision with a modest reduction in recall, yielding a more favorable precision-recall trade-off and broader recovery of curated regulatory interactions. Statistical evaluation revealed smaller differences between methods (Fig. 3A, 3B, bottom). GRASP showed slightly higher false omission rates and lower Wasserstein distances than GRNBoost. This gap is expected: statistical metrics primarily reflect perturbation-induced distributional shifts, which require expression-level data that LLM-based methods intentionally forgo. Nevertheless, GRASP remained competitive with GRNBoost across statistical metrics and experimental regimes, despite operating without any expression input.

**Figure 3.**
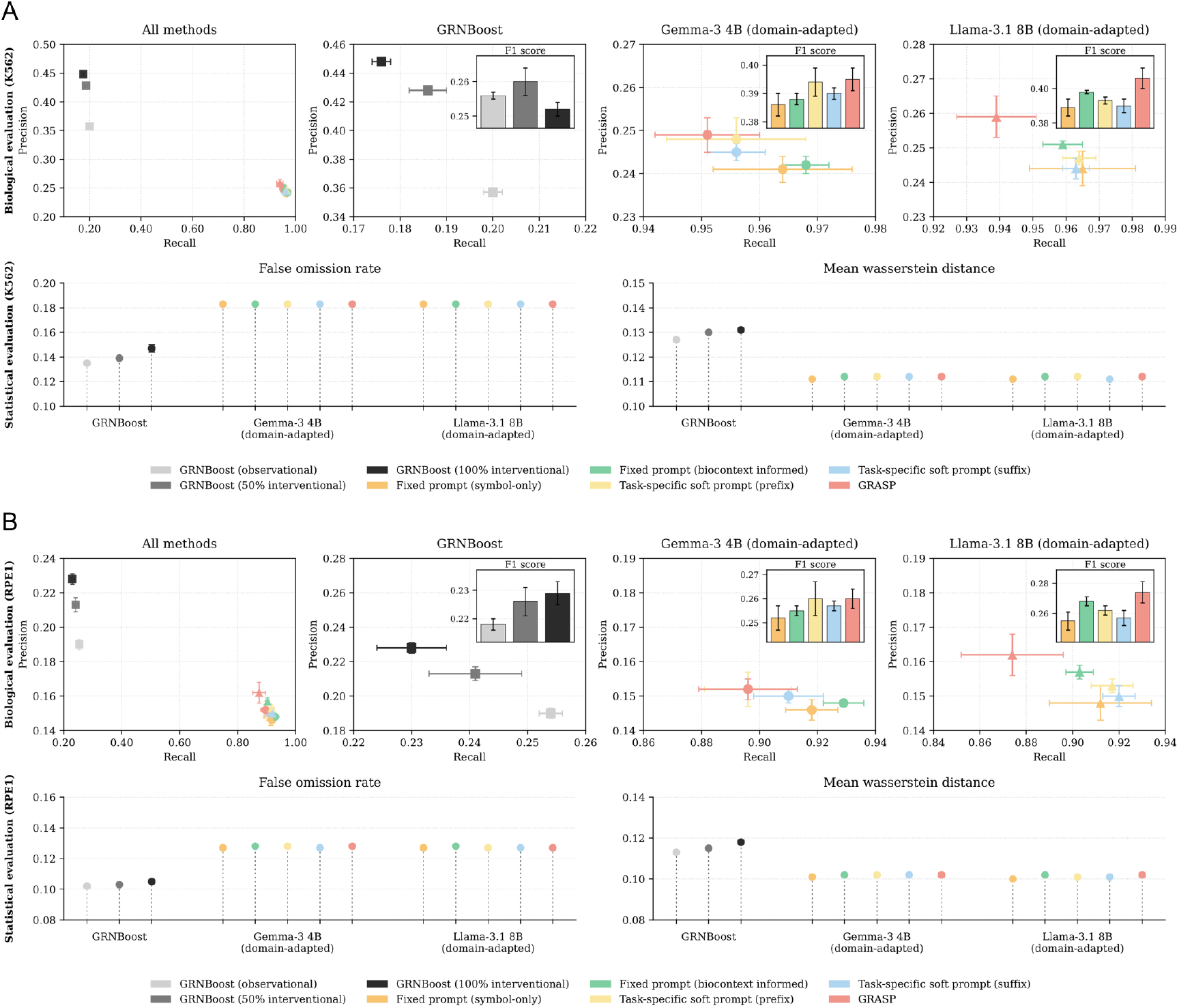
Biological and statistical evaluation on CausalBench. **(A)** Results for the K562 cell line. Top: precision-recall performance under biological evaluation, comparing GRNBoost and LLM-based methods (F1 scores shown in insets). Bottom: statistical evaluation, including false omission rate (left) and mean Wasserstein distance (right). **(B)** Results for the RPE1 cell line, shown in the same format.

Together, these results suggest that GRASP captures generalizable biological relationships that transfer from physical interaction data to regulatory contexts. Its strong biological performance, achieved without access to expression data, highlights the complementary value of language-based representations for gene network inference.

### GRASP achieves top performance in phosphorylation network inference

We next evaluated GRASP on a distinct class of gene interactions by applying the framework to phosphorylation networks, focusing on kinase-substrate relationships. Using the PhosphoNetworks^29^ database, a balanced dataset of 8,750 kinase-substrate pairs was constructed. The task was formulated as undirected interaction prediction to assess the model’s ability to identify general functional associations. Compared with previous sections, this dataset is substantially smaller and thus provides a low-data setting for evaluation. It was split into training (5,600 samples), validation (1,400 samples) and test (1,750 samples) sets. Training settings and evaluation metrics followed those used in the PPI experiments.

GRASP consistently outperformed all baselines in this setting (Fig. 4A). In precision-recall analysis, few-shot inference clustered well below the high-performing region, while GRASP occupied the upper-right corner with several points of precision gain over fixed prompts and task-specific soft prompts at comparable recall. ROC-AUC analyses confirmed these trends, with GRASP achieving the highest scores across all configurations (Gemma-3 4B: 0.953-0.963; Llama-3.1 8B: 0.960-0.965), and domain adaptation providing additional gains (Fig. 4B). Accuracy results were consistent with these findings (Supplementary Fig. 1C).

**Figure 4.**
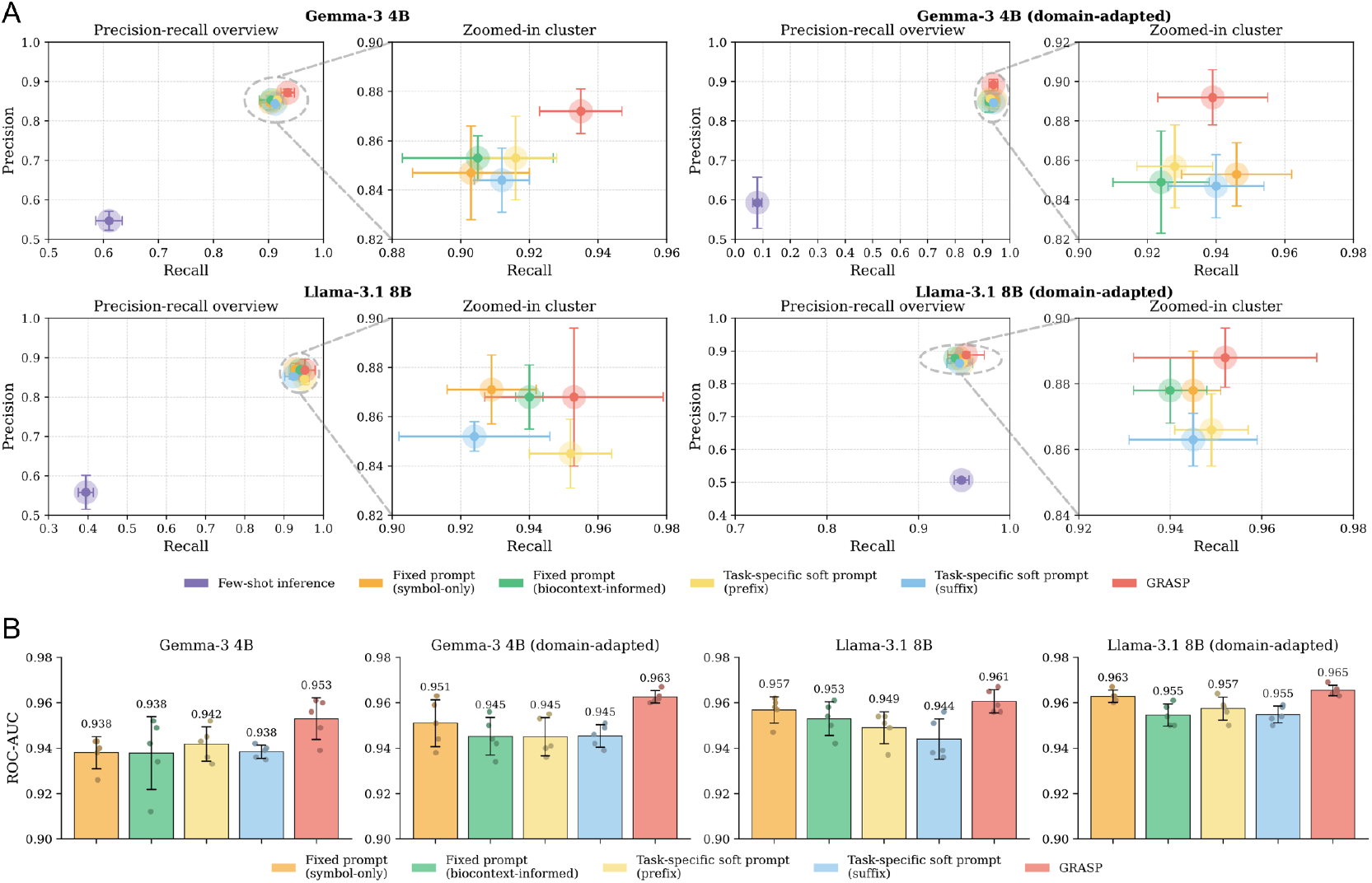
Comparative performance on the kinase-substrate relationship inference. **(A)** Precision-recall analyses for Gemma-3 4B and Llama-3.1 8B models before and after domain adaptation. Each panel shows an overview plot (left) and a zoomed-in view of high-performing clusters (right). **(B)** ROC-AUC scores for all prompting methods.

A notable observation is that augmenting fixed prompts with biological gene descriptions actually reduced performance relative to symbol-only prompts. This counterintuitive result suggests that generic textual descriptions can introduce noise when the prediction task involves a specialized interaction type such as phosphorylation, where only a narrow subset of gene function is relevant. GRASP avoids this pitfall by compressing gene summaries into compact latent vectors and synthesizing prompt tokens through a task-adapted factorized module, rather than directly appending verbose text to the input.

### GRASP recovers unannotated gene interactions from synthetic negative sets

A key question beyond benchmark performance is whether a model can identify genuine biological interactions that are absent from current annotations. Although our human PPI benchmark incorporated over one million experimentally validated pairs from multiple resources, the negative sets are constructed by random sampling, and some synthetic negatives inevitably correspond to real but unannotated interactions. An effective inference framework should assign these hidden positives higher scores than true negatives, reflecting a capacity for biological discovery beyond curated databases.

To test this, we cross-referenced our synthetic negative set (1,063,562 pairs) against the Integrated Interactions Database (IID)^27^, which contains 561,029 independently verified human PPIs. This identified 164 gene pairs that were labeled as negatives in our benchmark but are experimentally validated interactions in IID. We compared prediction scores for these 164 hidden positives against 10,000 randomly sampled true negatives, with no overlap between sets. Separation between the two score distributions was quantified using Wasserstein distance, which captures global distributional divergence, and Cohen’s d, which measures standardized effect size.

Across all configurations, GRASP achieved the largest separation between hidden positives and true negatives (Fig. 5A). With domain-adapted Llama-3.1 8B, GRASP attained a Wasserstein distance of 0.463 (±0.024) and a Cohen’s d of 1.880 (±0.051). Cohen’s d score indicated near-complete distributional separation between the two groups. By comparison, fixed prompts and task-specific soft prompts showed substantially smaller effect sizes. These results demonstrate that GRASP not only fits training labels accurately but also captures underlying biological signals that extend beyond the annotated dataset, effectively distinguishing genuine interactions even when they are mislabeled as negatives during training.

**Figure 5.**
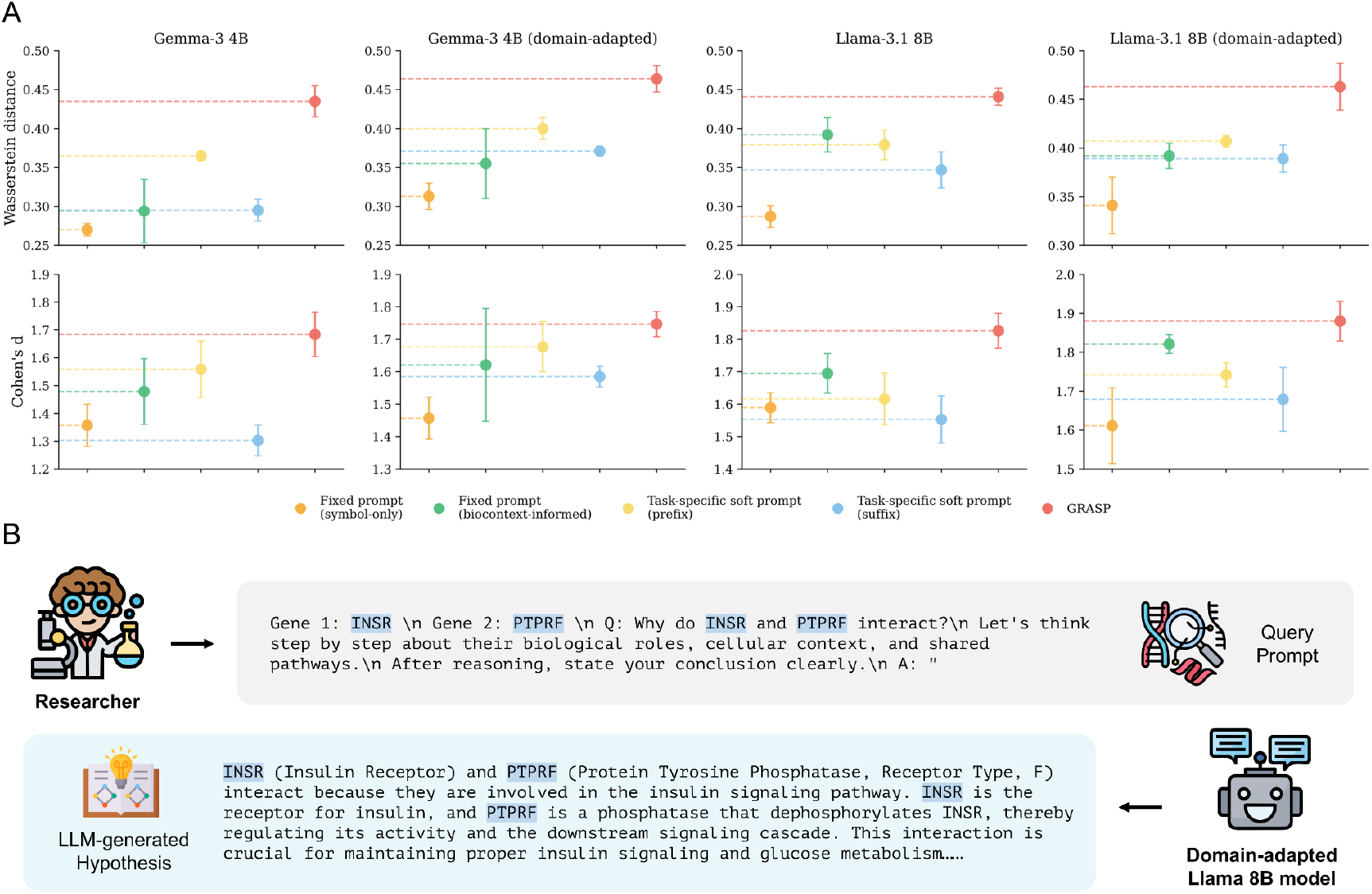
Recovery of previously unannotated gene interactions. **(A)** Separation between unannotated positives and true negatives measured by Wasserstein distance and Cohen’s d. **(B)** Example of hypothesis generation for a recovered interaction using the domain-adapted Llama-3.1-8B model. For the gene pair INSR-PTPRF, the model receives a zero-shot chain-of-thought style prompt and produces a reasoning-based explanation.

To illustrate the biological plausibility of these recovered interactions, we queried the domain-adapted Llama-3.1 8B model with a zero-shot chain-of-thought prompt^22^ (Fig. 5B). For the candidate interaction between INSR and PTPRF, the model generated a reasoning-based explanation identifying their shared involvement in insulin signaling: INSR as the insulin receptor and PTPRF as a phosphatase that modulates INSR phosphorylation and downstream signaling. This hypothesis is consistent with prior studies demonstrating that PTPRF dephosphorylates the insulin receptor and attenuates insulin signaling in cellular and animal models^30,31^. While such LLM-generated hypotheses often align with established knowledge, novel predictions ultimately require experimental validation. Additional representative examples are provided in Supplementary Table 1.

## Discussion

In this work, we introduce GRASP, an instance-adaptive prompting framework for LLM-based gene network inference. Unlike task-specific soft prompts that apply a single shared embedding to all inputs, GRASP first reasons about each gene individually by generating a concise summary, and this gene-level knowledge is then compressed into a fixed vector that conditions pairwise inference. This two-stage design decouples gene-level knowledge extraction from interaction prediction, allowing each stage to serve a distinct role: the first externalizes what the model knows about each gene, and the second distills that knowledge into a compact, task-relevant signal for prompt synthesis.

Across the three benchmark tasks on distinct gene network types: protein-protein interactions, single-cell perturbation-derived regulatory relationships, and kinase-substrate phosphorylation networks, GRASP consistently outperformed all prompting baselines. Improvements in ROC-AUC across settings suggest that GRASP enhances the ranking of candidate interactions rather than merely shifting decision thresholds, a property particularly valuable for practical applications where prioritizing high-confidence predictions matters more than binary classification.

The cross-species transfer results reveal an interesting exception. While GRASP achieved the highest ROC-AUC in most cross-species conditions, the overall performance gap between methods was narrower than in the within-species setting. We attribute this to the experimental design: all gene pairs overlapping with the human training set were removed from the cross-species benchmarks, leaving only interactions that are not shared across species. These remaining pairs likely represent species-specific interactions that lack strong representation in the LLM’s text-derived knowledge, making them inherently more difficult for any language-based method. Even in such a scenario, GRASP still maintains a ranking advantage, suggesting it captures transferable relational patterns beyond simple gene-name recognition.

The most distinctive finding is GRASP’s capacity to recover potentially unannotated interactions. The near-complete distributional separation between hidden positives and true negatives (Cohen’s d ≈ 1.88) indicates that GRASP does not simply memorize the knowledge inside curated databases but learn and capture biological signals from them. This result has practical implications: high-confidence GRASP predictions could serve as a prioritization tool for experimental validation, complementing existing interaction databases with computationally nominated candidates. The LLM-generated hypotheses for recovered interactions, such as the INSR-PTPRF example grounded in insulin signaling, further suggest that the model has internalized mechanistic knowledge that goes beyond co-occurrence statistics in the training corpus.

The consistent advantage of GRASP over both task-specific soft prompts and biocontext-informed fixed prompts provides insight into why the framework is effective. Incorporating biological context by appending gene descriptions directly to the input prompt did not consistently improve performance, as natural language descriptions often include redundant details that can obscure subtle relational signals. In the phosphorylation network inference task, for example, prompts with additional gene descriptions degraded accuracy relative to symbol-only prompts. GRASP addresses this by encoding gene summaries into fixed-length latent vectors before prompt synthesis, implicitly filtering heterogeneous information and enabling efficient, adaptive conditioning through only three soft prompt tokens.

Despite these strengths, several limitations remain. First, GRASP inherits the knowledge biases of its underlying LLM: well-studied genes with extensive literature are likely better represented than poorly characterized ones, potentially skewing predictions toward known biology. Stratifying performance by gene-level publication count could help quantify this bias. Second, the current framework models all interactions as undirected binary relationships and does not capture directionality. Extending GRASP to directed or hypergraph formulations would broaden its applicability to a wider range of biological networks. Third, GRASP operates entirely on text-derived representations without incorporating experimental data such as gene expression profiles or protein structures. While this design enables generalization across interaction types, developing hybrid frameworks that fuse adaptive prompting with expression-based or structure-based features could improve performance on tasks where textual knowledge alone is insufficient.

Overall, GRASP establishes that instance-adaptive prompt conditioning is a viable and effective strategy for LLM-based biological prediction. As LLMs continue to internalize an expanding body of biomedical knowledge, frameworks that can efficiently and adaptively query this knowledge will become increasingly valuable for biological discovery.

## Methods

### Human protein-protein interaction dataset construction

We collected experimentally validated human physical protein-protein interaction (PPI) pairs from four curated databases: BioGRID^23^, MIPS^24^, IntAct^25^, and STRING^26^. In total, 1,063,562 gene pairs were obtained and treated as positive pairs. To focus on language-based inference, we retained only the gene symbols and discarded all other associated information.

To construct a balanced dataset, synthetic negative pairs were generated by sampling gene pairs absent from all curated PPI sources. Assuming undirected interactions, we first extracted unique gene symbols from the positive set and enumerated all possible gene-gene combinations. Negative pairs were randomly sampled from these combinations, excluding any overlapping with the positives. This sampling process continues until the number of negative samples matches the number of positive ones. Following this procedure, we retrieved a total of 2,127,124 gene pairs, with equal numbers of positive and negative samples.

### Cross-species protein-protein interaction dataset construction

Cross-species datasets for chicken, cow and dog were constructed using physical PPI pairs from the Integrated Interactions Database (IID)^27^. IID integrates multiple types of evidence, including experimental detection, orthology, and machine learning predictions. To ensure high-quality data, we retained only experimentally verified PPIs. We further removed any pairs overlapping with the human PPI dataset used in earlier experiments. Balanced datasets were constructed by generating an equal number of negative gene pairs using the same procedure as above, resulting in 153,252 pairs for chicken, 165,574 for cow, and 149,960 for dog.

### Phosphorylation network dataset construction

We derived phosphorylation relationships from a kinase-substrate relationship (KSR) study^29^. Gene pairs in the combined KSR dataset, including 3,656 refined KSRs and 744 literature-curated KSRs, were treated as positive samples. An equal number of negative pairs (4,375) was generated using the same sampling procedure as above. To ensure the validity of the negative set, pairs appearing in the raw KSR dataset (24,046 KSRs identified through protein microarray assays) were excluded. The final kinase-substrate interaction dataset comprises 8,750 pairs.

### Gene summary curation for BioContext-informed prompts

To construct BioContext-informed prompts, we retrieved gene summaries from MyGene.info (https://mygene.info)^32^. Gene symbols were queried in bulk using the “querymany” function from the Mygene Python package (scopes = “symbol”, fields = “summary”). The returned summaries were parsed to extract only the first sentence for each gene to accommodate input token limitations. We incorporate these summaries into BioContext-informed prompts alongside gene symbols.

### Gene-relation adaptive soft prompt

GRASP is a gene-relation adaptive soft prompt tuning method that appends a small number of learnable virtual tokens conditioned on a queried gene pair.

#### 1. Gene vector encoding

Let the backbone language model have hidden size *d*. For each gene *g*, we derive a fixed vector representation *s*_*g*_ ∈ ℝ^*d*^ that captures its biological context. To obtain *s*_*g*_, we first prompt the LLM to generate a short textual summary of gene *g*. The summary is then encoded using the same backbone model, and the final-layer hidden states are mean-pooled over the token dimension to produce a single *d* -dimensional vector. All gene representations are precomputed and stored in a look-up table *S* and remain fixed during downstream fine-tuning.

#### 2. Factorized soft prompt synthesis

The core idea of GRASP is to generate a soft prompt embedding from a context vector *z* ∈ ℝ^*d*^ using a factorized formulation. Specifically, GRASP defines a mapping

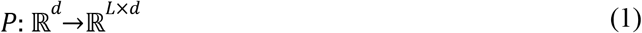

where *L* denotes the prompt length. For a given context vector *z*, the prompt embedding is defined as:

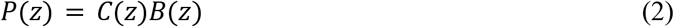

*C*(*z*) ∈ ℝ^*L*×*r*^ is a gene-specific coefficient matrix and *B*(*z*) ∈ ℝ^*r*×*d*^ is a shared gene prototype matrix.

The coefficient matrix *C*(*z*) is obtained via a linear projection:

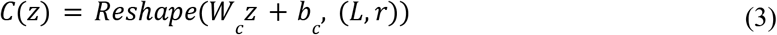

where *W*_*c*_ ∈ ℝ ^(*Lr*)×*d*^ and *b*_*c*_ ∈ ℝ ^*Lr*^.

The prototype matrix *B*(*z*) is defined as a convex combination of *K* global basis matrices 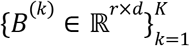:

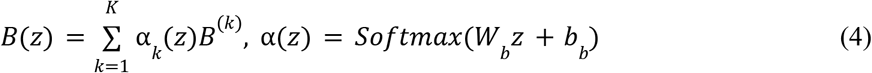

where α_*k*_ (*z*) ∈ ℝ^*K*^, α_*k*_ (*z*) ≥ 0 and 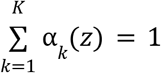. This factorization allows prompt embeddings to adapt to each input through *C*(*z*), while maintaining shared structure via the global basis bank in *B*(*z*). The internal rank *r* controls representational capacity, and the number of bases *K* determines the diversity of the shared prototype space.

#### 3. Gene soft prompt construction

Given a gene pair (*a, b*), GRASP constructs three prompt segments. Two gene-specific segments are generated using the same parameterization with inputs *z* = *s*_*a*_ and *z* = *s*_*b*_, respectively.

The third segment encodes the relationship between the two genes. Its context vector is defined as:

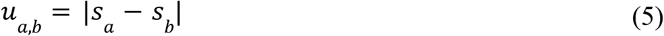

which captures contrastive biological features. The relational prompt is generated using a separate set of projection parameters, while sharing the same global basis bank:

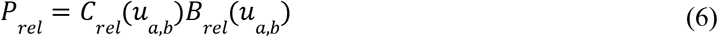

The final prompt for gene pair (*a, b*) is obtained by concatenation:

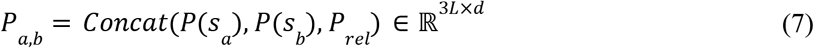

This prompt is appended as a suffix to the input embeddings. With *L* = 1, GRASP introduces exactly three adaptive virtual tokens per example.

#### 4. Interpretation and efficiency

This factorized formulation clarifies how GRASP balances instance-specific adaptation and parameter sharing. The coefficient matrix *C*(*z*) captures gene-specific information by selecting directions within a low-dimensional subspace, while the prototype matrix *B*(*z*) provides shared structure through a small set of global bases. This structure stabilizes training and promotes generalization across semantically related genes.

Despite its compact design, GRASP remains expressive. Even with *L* = 1, each adaptive token is derived from a full *r* × *d* prototype matrix, enabling rich context-dependent modulation. At the same time, efficiency is maintained through lightweight projections and a small number of learnable parameters.

### Classification head and training objective

To map token-level representations to an interaction probability, we attach a classification head to the frozen LLM backbone, replacing the original language modeling head. Let *X* ∈ ℝ^*T*×*d*^ denote the final-layer hidden states for an input sequence of length *T*.

A position-wise three-layer MLP is applied independently to each token embedding to produce scalar local logits:

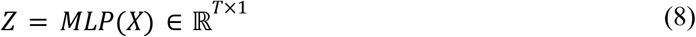

where

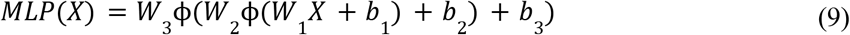

and ϕ(·) denotes the SiLU^33^ activation function.

The token-level logits are aggregated into a sequence-level logit using a learnable linear pooling:

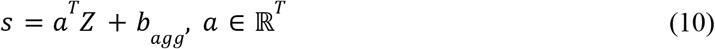

where *a* contains position-specific weights and *b*_*agg*_ is a scalar bias.

The predicted interaction probability is:

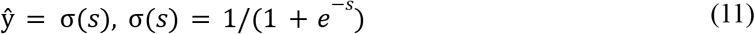

Given a binary label *y* ∈ {0, 1}, the model is trained with the batch-averaged binary cross-entropy (BCE) loss:

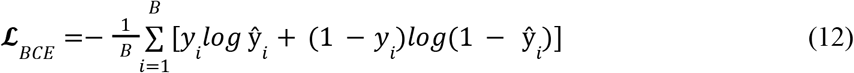

where *B* denotes the batch size.

During fine-tuning, only the classification head and soft prompt parameters are updated, while the backbone model remains frozen.

### Hyperparameter settings

Across all experiments, we evaluated Gemma-3-4B-IT^19^ and Llama-3.1-8B-Instruct^20^ models, together with their domain-adapted variants pretrained on biomedical corpora. All fine-tuning runs were conducted for 15 epochs. Training used bfloat16 (BF16) precision and the AdamW optimizer with default settings from Hugging Face Transformers, including a weight decay of 0.01, a learning rate of 1 × 10^−4^, and a warmup ratio of 0.05. The batch size was set to 128 for PPI network experiments and 64 for phosphorylation network experiments.

#### Few-shot inference

Models were prompted with six in-context examples (three positive and three negative gene-pair relations), followed by a query pair. Responses were generated with a maximum sequence length of 512 tokens, and predictions were extracted by parsing the generated text.

#### Fix prompt

Two prompt variants were evaluated: (i) Symbol-only prompts with a sequence length of 64 tokens, and (ii) BioContext-informed prompts incorporating brief gene summaries, with a sequence length of 128 tokens.

#### Task-specific soft prompt

The original query was tokenized to a maximum length of 80 tokens using left padding with the EOS token. Three task-specific soft prompt tokens were concatenated to the input query as either a prefix or a suffix, resulting in a total input length of 83 tokens.

#### Gene-relation adaptive soft prompt

GRASP generated three adaptive soft prompt tokens per gene pair, comprising two gene-specific tokens and one relation-specific token. These tokens were appended as a suffix to the input sequence, while all other preprocessing steps remained the same as in the task-specific soft prompt setting. The internal rank r was set to 1024 for PPI network experiments and 512 for phosphorylation network experiments. In both settings, K = 4 prototype bases were used. Gene vectors were constructed from LLM-generated one-paragraph gene summaries, with up to 128 tokens per summary.

## Data availability

Human physical protein-protein interactions were collected from publicly available resources, including BioGRID (https://downloads.thebiogrid.org/BioGRID), MIPS (http://mips.gsf.de/proj/ppi/), IntAct (https://www.ebi.ac.uk/intact/download), and STRING (https://string-db.org/cgi/download.pl). Cross-species protein-protein interactions for chicken, cow and dog were obtained from Integrated Interactions Database (IID) (https://iid.ophid.utoronto.ca/). Single-cell perturbation datasets in the CausalBench benchmark were automatically downloaded during evaluation. The corresponding dataset URLs are specified in the causalscbench/data_access/datasets/download_evaluation_files.py script within the GitHub repository (https://github.com/causalbench/causalbench). Kinase-substrate relationship dataset was obtained from the PhosphoNetworks (https://phosphonetworks.org/download.html).

## Code availability

The source code is publicly available at https://github.com/GuanLab/GRASP.

## Reference

1. Badia-i-Mompel, P. et al. Gene regulatory network inference in the era of single-cell multi-omics. Nat. Rev. Genet. 24, 739–754 (2023).

2. Xia, S., Chen, J., Arsala, D., Emerson, J. J. & Long, M. Functional innovation through new genes as a general evolutionary process. Nat. Genet. 1–15 (2025).

3. Bonev, B. et al. Opportunities and challenges of single-cell and spatially resolved genomics methods for neuroscience discovery. Nat. Neurosci. 27, 2292–2309 (2024).

4. Zhu, Q. M. et al. Protein interaction networks in the vasculature prioritize genes and pathways underlying coronary artery disease. Commun. Biol. 7, 87 (2024).

5. Kim, S. & Wysocka, J. Deciphering the multi-scale, quantitative cis-regulatory code. Mol. Cell 83, 373–392 (2023).

6. Sevimoglu, T. & Arga, K. Y. The role of protein interaction networks in systems biomedicine. Comput. Struct. Biotechnol. J. 11, 22–27 (2014).

7. Mercatelli, D., Scalambra, L., Triboli, L., Ray, F. & Giorgi, F. M. Gene regulatory network inference resources: A practical overview. Biochim. Biophys. Acta BBA-Gene Regul. Mech. 1863, 194430 (2020).

8. De Oliveira, P. S. L. et al. Revisiting protein kinase–substrate interactions: Toward therapeutic development. Sci. Signal. 9, (2016).

9. Sledzieski, S., Singh, R., Cowen, L. & Berger, B. D-SCRIPT translates genome to phenome with sequence-based, structure-aware, genome-scale predictions of protein-protein interactions. Cell Syst. 12, 969–982.e6 (2021).

10. Hashemifar, S., Neyshabur, B., Khan, A. A. & Xu, J. Predicting protein–protein interactions through sequence-based deep learning. Bioinformatics 34, i802–i810 (2018).

11. Ko, Y. S., Parkinson, J., Liu, C. & Wang, W. TUnA: an uncertainty-aware transformer model for sequence-based protein–protein interaction prediction. Brief. Bioinform. 25, bbae359 (2024).

12. Aibar, S. et al. SCENIC: single-cell regulatory network inference and clustering. Nat. Methods 14, 1083–1086 (2017).

13. Brouillard, P., Lachapelle, S., Lacoste, A., Lacoste-Julien, S. & Drouin, A. Differentiable causal discovery from interventional data. Adv. Neural Inf. Process. Syst. 33, 21865–21877 (2020).

14. Nováček, V. et al. Accurate prediction of kinase-substrate networks using knowledge graphs. PLoS Comput. Biol. 16, e1007578 (2020).

15. Jha, K., Saha, S. & Singh, H. Prediction of protein–protein interaction using graph neural networks. Sci. Rep. 12, 8360 (2022).

16. Liévin, V., Hother, C. E., Motzfeldt, A. G. & Winther, O. Can large language models reason about medical questions? Patterns 5, (2024).

17. Wang, L. et al. Prompt engineering in consistency and reliability with the evidence-based guideline for LLMs. NPJ Digit. Med. 7, 41 (2024).

18. Lester, B., Al-Rfou, R. & Constant, N. The Power of Scale for Parameter-Efficient Prompt Tuning. Preprint at 10.48550/arXiv.2104.08691 (2021).

19. Team, G. et al. Gemma 3 Technical Report. Preprint at 10.48550/arXiv.2503.19786 (2025).

20. Grattafiori, A. et al. The llama 3 herd of models. ArXiv Prepr. ArXiv240721783 https://arxiv.org/abs/2407.21783 (2024).

21. Gururangan, S. et al. Don’t Stop Pretraining: Adapt Language Models to Domains and Tasks. Preprint at 10.48550/arXiv.2004.10964 (2020).

22. Kojima, T., Gu, S. S., Reid, M., Matsuo, Y. & Iwasawa, Y. Large language models are zero-shot reasoners. Adv. Neural Inf. Process. Syst. 35, 22199–22213 (2022).

23. Oughtred, R. et al. The BIOGRID database: A comprehensive biomedical resource of curated protein, genetic, and chemical interactions. Protein Sci. 30, 187–200 (2021).

24. Mewes, H.-W. et al. MIPS: a database for genomes and protein sequences. Nucleic Acids Res. 30, 31–34 (2002).

25. Del Toro, N. et al. The IntAct database: efficient access to fine-grained molecular interaction data. Nucleic Acids Res. 50, D648–D653 (2022).

26. Szklarczyk, D. et al. The STRING database in 2025: protein networks with directionality of regulation. Nucleic Acids Res. 53, D730–D737 (2025).

27. Kotlyar, M. et al. IID 2021: towards context-specific protein interaction analyses by increased coverage, enhanced annotation and enrichment analysis. Nucleic Acids Res. 50, D640–D647 (2022).

28. Chevalley, M., Roohani, Y. H., Mehrjou, A., Leskovec, J. & Schwab, P. A large-scale benchmark for network inference from single-cell perturbation data. Commun. Biol. 8, 412 (2025).

29. Hu, J. et al. Phospho Networks: a database for human phosphorylation networks. Bioinformatics 30, 141–142 (2014).

30. Zabolotny, J. M. et al. Overexpression of the LAR (leukocyte antigen-related) protein-tyrosine phosphatase in muscle causes insulin resistance. Proc. Natl. Acad. Sci. 98, 5187–5192 (2001).

31. Sevillano, J., Sánchez-Alonso, M. G., Pizarro-Delgado, J. & Ramos-Álvarez, M. del P. Role of receptor protein tyrosine phosphatases (RPTPs) in insulin signaling and secretion. Int. J. Mol. Sci. 22, 5812 (2021).

32. Wu, C., MacLeod, I. & Su, A. I. Biogps and MyGene. info: organizing online, gene-centric information. Nucleic Acids Res. 41, D561–D565 (2013).

33. Elfwing, S., Uchibe, E. & Doya, K. Sigmoid-weighted linear units for neural network function approximation in reinforcement learning. Neural Netw. 107, 3–11 (2018).

