## Supplementary Information for "GRASP: Gene-relation adaptive soft prompt for scalable and generalizable gene network inference with large language models"

**Supplementary Figure 1: Accuracy across species and interaction types. (A)** Performance on human PPI prediction using all methods (top) and excluding few-shot inference (bottom). **(B)** Cross-species transfer evaluation on chicken, cow and dog PPI datasets using models fine-tuned on human PPIs. **(C)** Performance on kinase-substrate relationship inference with all methods (top) and excluding few-shot inference (bottom).

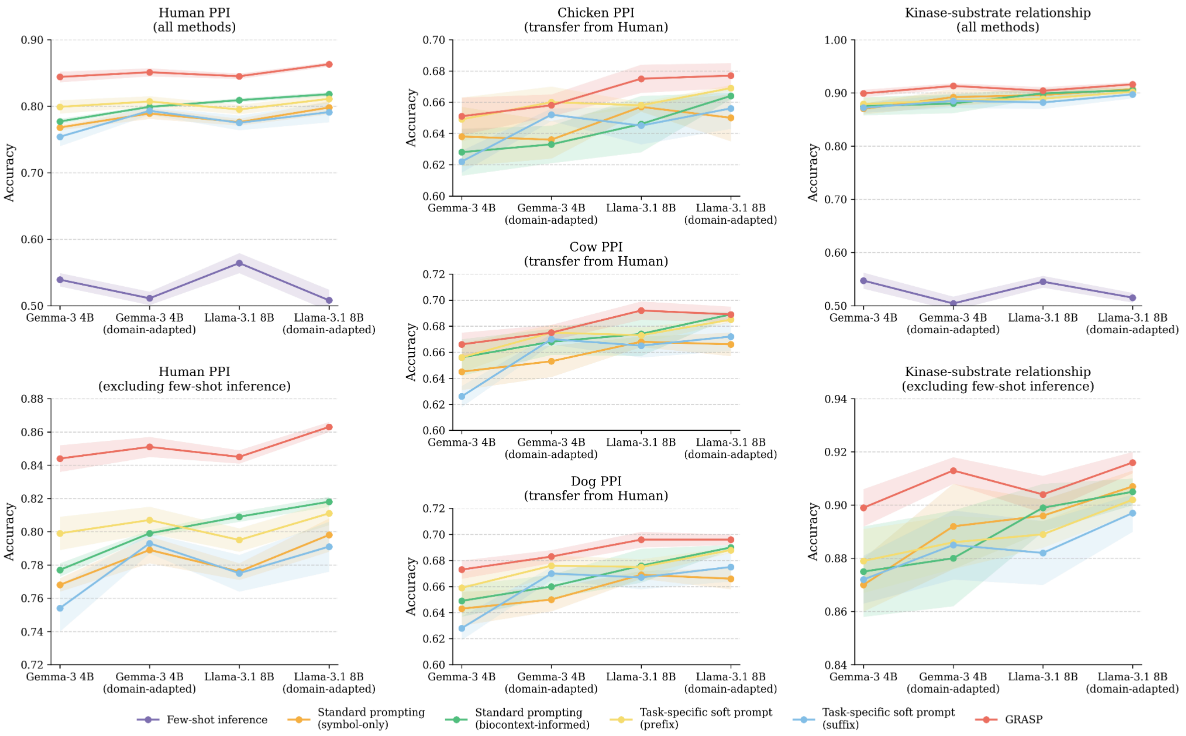


**Supplementary Figure 2: Per-gene error rate analysis. (A)** Scatter plots of per-gene error rate versus positive-interaction degree under five prompting strategies. Each gray dot represents a gene, with selected examples highlighted (COG4, SLC22A9, ZIC1, and CDCA7L). **(B)** Pairwise comparisons between GRASP (E) and other methods (A-D). The Δ error rate is plotted against positive-interaction degree (GRASP minus the comparison method). Points are colored by gene density, with blue lines denoting the mean Δ error rate per degree bin. Negative values indicate genes for which GRASP outperforms the comparison method.

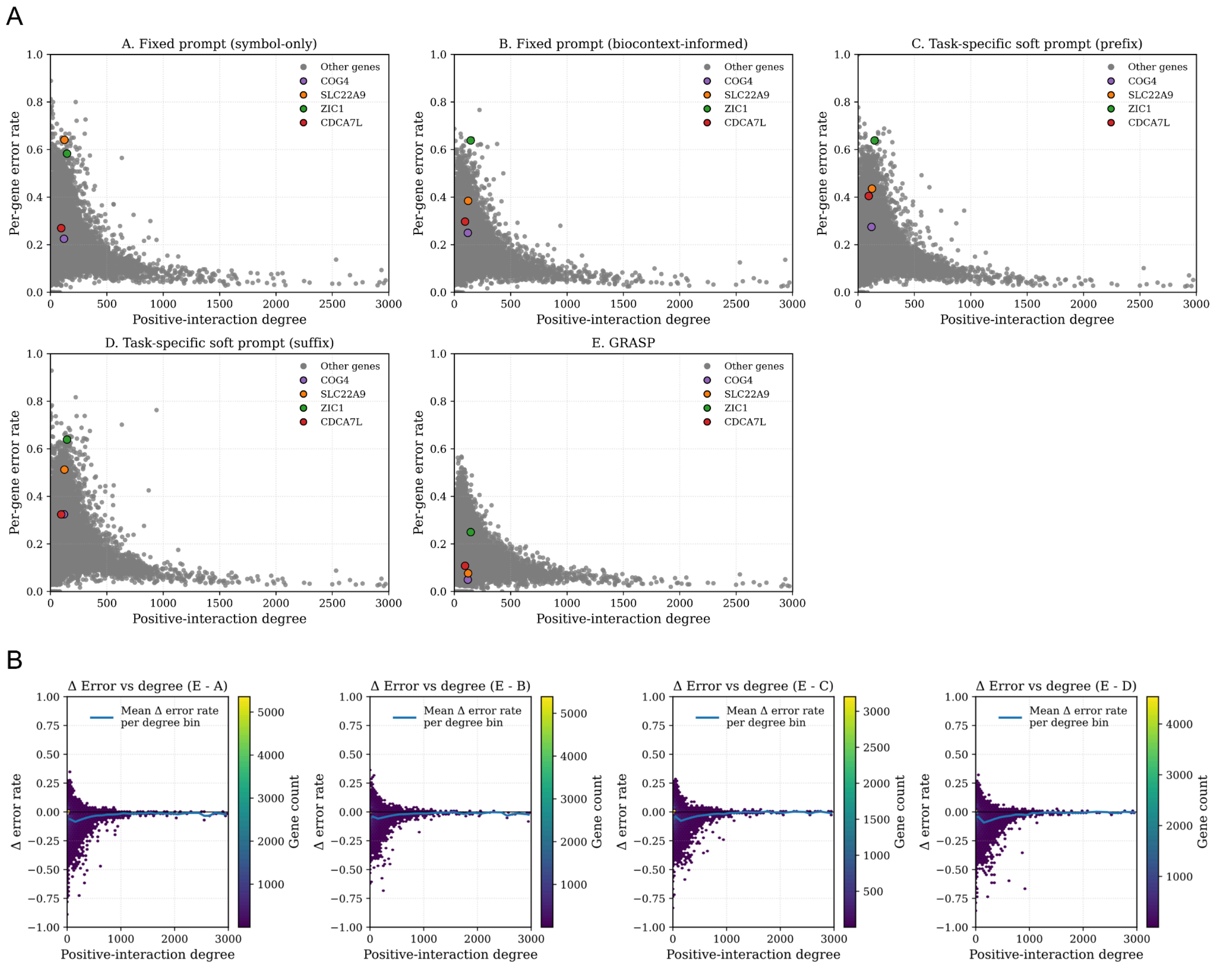
A
A. Fixed prompt (symbol-only)
B. Fixed prompt (biocontext-informed)
C. Task-specific soft prompt (prefix)
D. Task-specific soft prompt (suffix)
E. GRASP
Other genes
COG4
SLC22A9
ZIC1
CDCA7L
Per-gene error rate
Positive-interaction degree
B
Δ Error vs degree (E - A)
Δ Error vs degree (E - B)
Δ Error vs degree (E - C)
Δ Error vs degree (E - D)
Mean Δ error rate per degree bin
Δ error rate
Gene count
Positive-interaction degree

**Supplementary Figure 3: Subnetwork visualizations of representative genes.** Subnetworks centered on four representative genes—COG4 (Golgi retrograde transport), SLC22A9 (hepatic anion transport), ZIC1 (cerebellar transcription), and CDCA7L (MYC-associated chromatin regulation)—are shown under different prompting strategies. Edge styles and colors follow the legend at the bottom: solid blue, true positives (TPs); solid red, false negatives (FNs); dashed red, false positives (FPs). True negatives (TNs) are omitted for clarity.

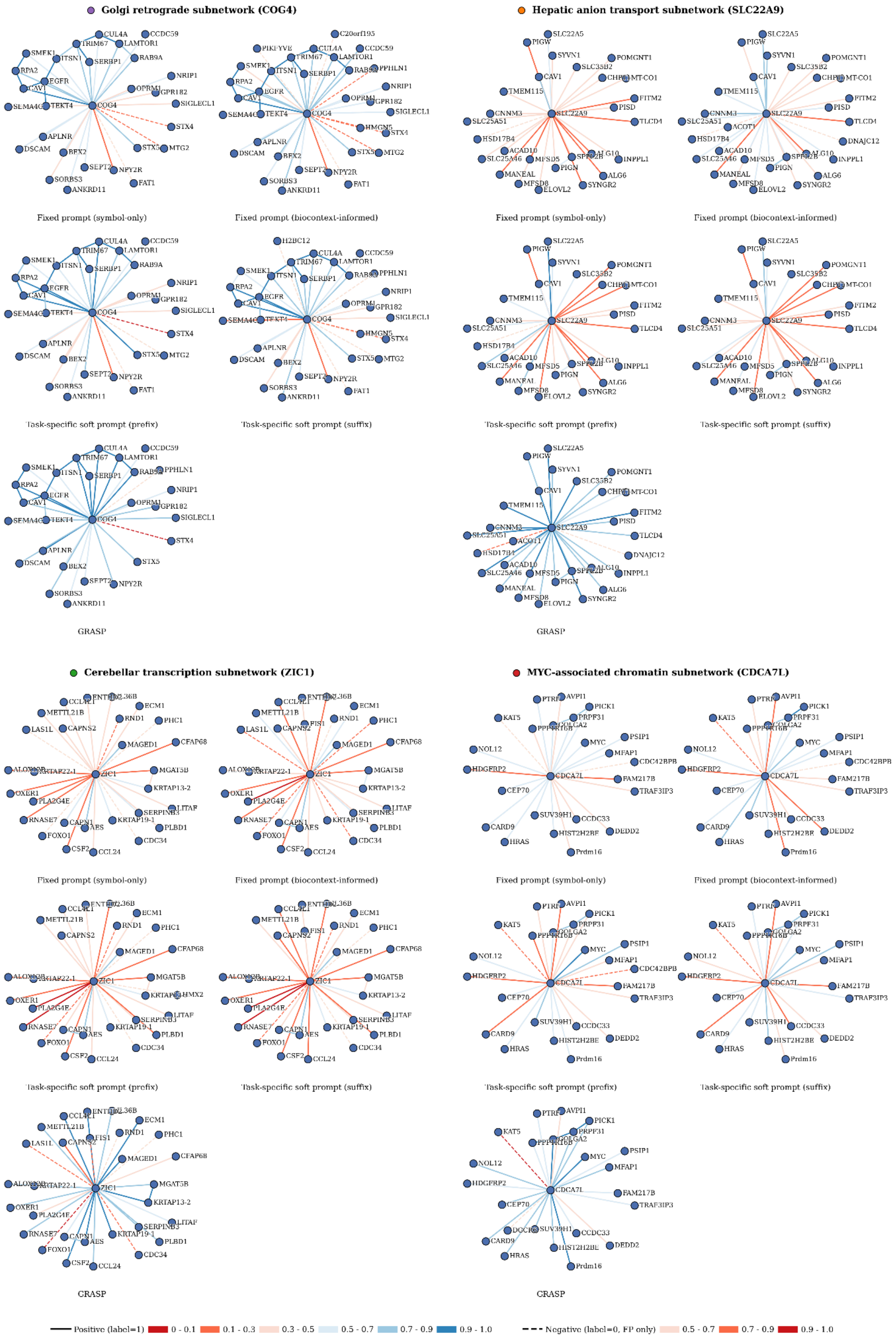

**Supplementary Figure 4: Cross-species transfer evaluation in PPI inference. (A)** Precision-recall analysis on chicken, cow and dog PPI datasets using models fine-tuned on human PPIs. To highlight performance differences, we overlaid F1-score contours at 0.02 intervals, increasing from the lower left to the upper right; methods positioned further toward the upper-right region therefore achieve higher F1 values. **(B)** ROC-AUC results for the same setting.

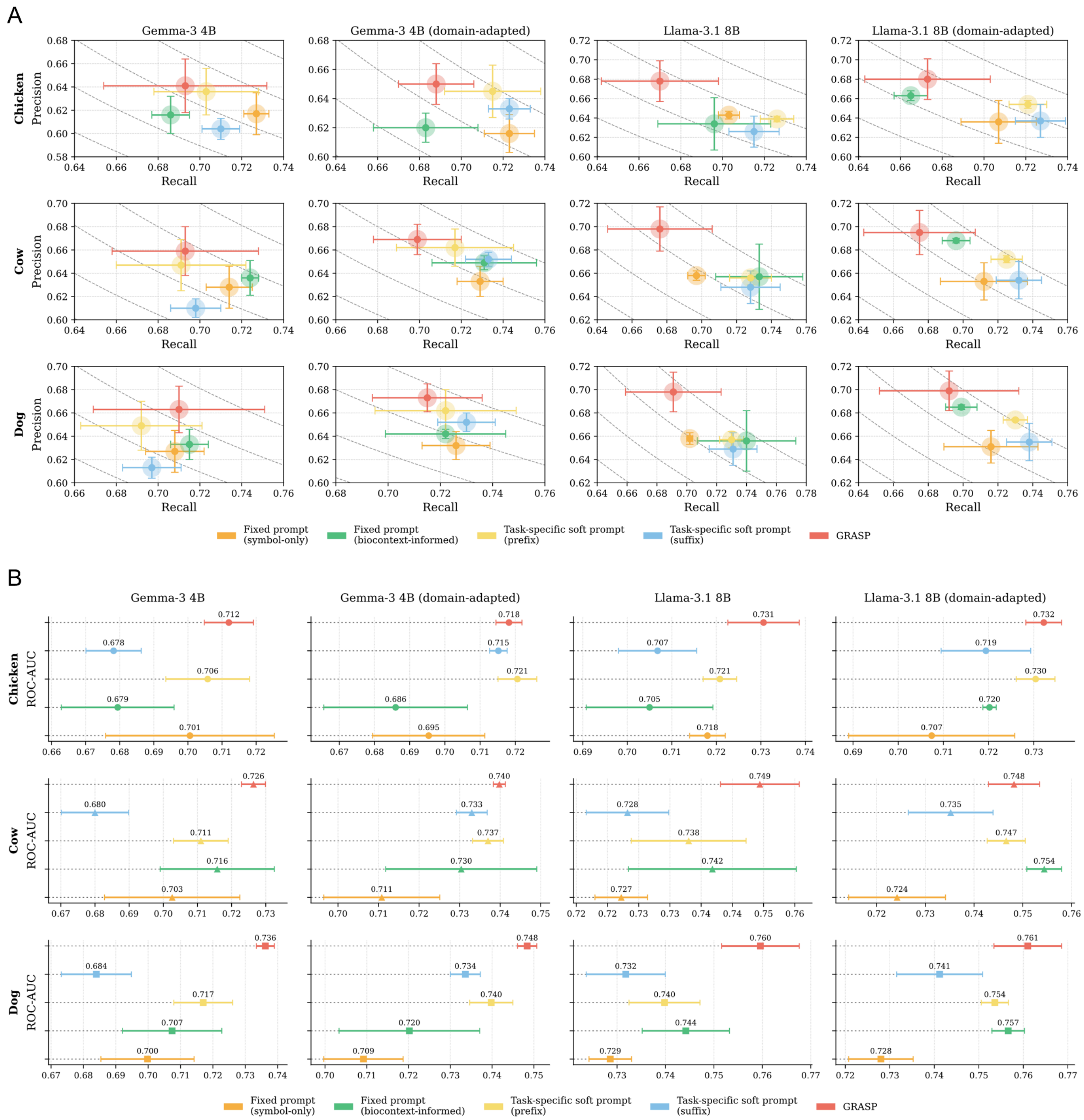

**Supplementary Table 1: Example LLM-generated hypotheses for gene interactions.** Five representative gene pairs are shown with their corresponding hypotheses generated by the domain-adapted Llama-3.1-8B model. For each pair, the model produces a mechanistic explanation of potential interactions.

| Gene A | Gene B | LLM-generated hypothesis |
|---|---|---|
| INSR | PTPRF | INSR (Insulin Receptor) and PTPRF (Protein Tyrosine Phosphatase, Receptor Type, F) interact because they are involved in the insulin signaling pathway. INSR is the receptor for insulin, and PTPRF is a phosphatase that dephosphorylates INSR, thereby regulating its activity and the downstream signaling cascade. This interaction is crucial for maintaining proper insulin signaling and glucose metabolism. \nThis interaction is an example of a regulatory mechanism in cellular signaling pathways, where a phosphatase (PTPRF) modulates the activity of a receptor (INSR) to control the flow of signals within the cell. \nThe interaction between INSR and PTPRF is a specific example of the complex interplay between different proteins in cellular signaling pathways, highlighting the importance of precise regulation in maintaining proper cellular function. |
| PAK1 | STMN1 | PAK1 and STMN1 interact because they are involved in the regulation of the actin cytoskeleton, a key process in cell migration, division, and signaling. PAK1 is a serine/threonine kinase that activates LIMK1, which in turn phosphorylates and inactivates cofilin, a protein that depolymerizes actin filaments. STMN1, on the other hand, is a microtubule-severing protein that also regulates the actin cytoskeleton. The interaction between PAK1 and STMN1 allows for the coordination of actin and microtubule dynamics, enabling cells to undergo shape changes and movement. This interaction is crucial for various cellular processes, including cell migration, division, and signaling. \nThis answer is based on the understanding of the biological roles of PAK1 and STMN1, their involvement in the regulation of the actin cytoskeleton, and their shared pathways. The interaction between these two proteins is essential for the proper functioning of cells, particularly in processes that require dynamic changes in the cytoskeleton. |
| PARP1 | TCF4 | PARP1 and TCF4 interact because they are involved in the regulation of gene expression and chromatin remodeling. PARP1 is a nuclear enzyme that plays a key role in DNA repair, transcriptional regulation, and chromatin remodeling. TCF4 is a transcription factor that regulates the expression of genes involved in cell growth, differentiation, and survival. Their interaction may be crucial for the regulation of gene expression and chromatin remodeling in response to DNA damage or other cellular stresses. This interaction may also be relevant in the context of neurodegenerative diseases, such as |

| | | |
|---|---|---|
| | | Down syndrome, where TCF4 is implicated, and cancer, where PARP1 is often overexpressed. \nThis interaction may be mediated by the ability of PARP1 to modify chromatin structure and recruit transcriptional regulators, including TCF4, to specific genomic regions. The interaction between PARP1 and TCF4 may also be influenced by the presence of other proteins or post-translational modifications that modulate their activity and localization. |
| H3C7 | YWHAB | H3C7 and YWHAB interact because they are both involved in the regulation of gene expression and cellular signaling pathways. H3C7 is a histone modification enzyme that plays a role in chromatin remodeling and gene regulation, while YWHAB is a 14-3-3 protein that binds to phosphoserine or phosphothreonine residues on target proteins, influencing their activity and localization. Their interaction likely facilitates the regulation of gene expression and cellular signaling pathways, particularly in response to cellular stress or changes in the cellular environment.\nThe final answer is: They interact to regulate gene expression and cellular signaling pathways. |
| ACTC1 | HSPB1 | ACTC1 and HSPB1 interact because they are both involved in maintaining cellular homeostasis and responding to stress. ACTC1 is a component of the sarcomere, the contractile unit of muscle cells, and plays a critical role in muscle contraction and relaxation. HSPB1, on the other hand, is a heat shock protein that acts as a molecular chaperone, helping to maintain protein structure and function under stress conditions. Given their roles in maintaining cellular integrity and responding to stress, it is likely that ACTC1 and HSPB1 interact to coordinate their activities and ensure proper muscle function under various conditions. |
